## Supplemental Figures for "Hetero-oligomerization of TDP-43 carboxy-terminal fragments with cellular proteins contributes to proteotoxicity"

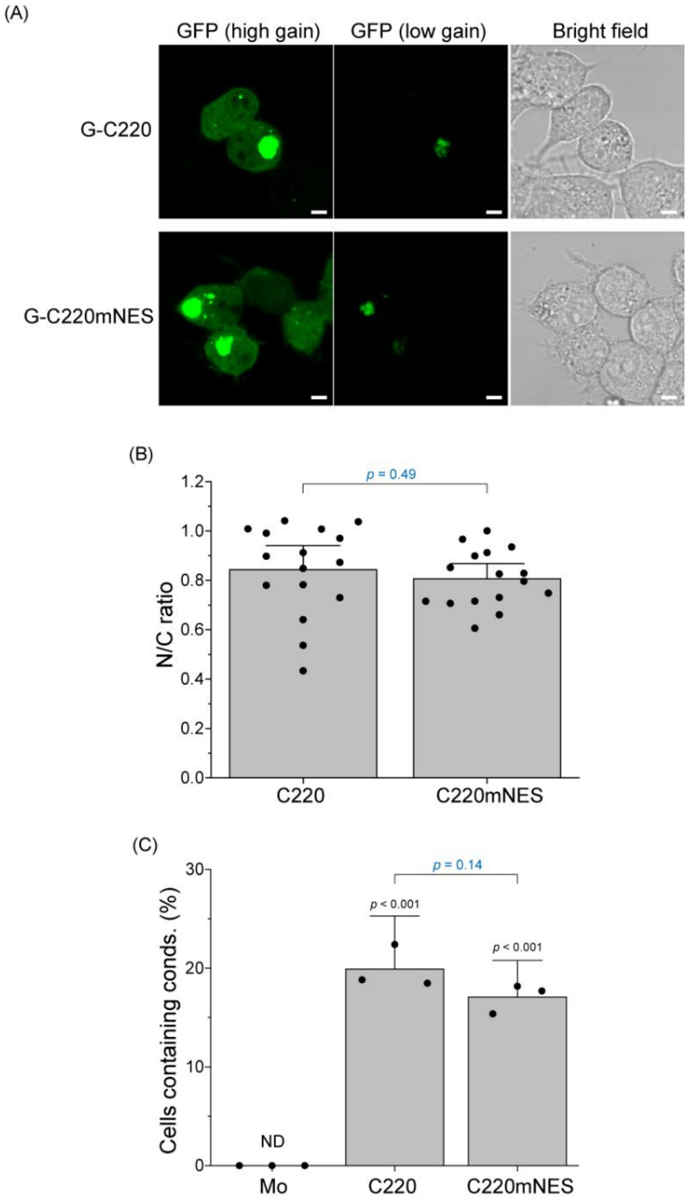


**Supplemental Figure 1: Subcellular distribution and cytoplasmic condensation of C220 carrying mutations for nuclear exporting signal sequence in Neuro2a cells**

(A) Confocal images of Neuro2a cells expressing GFP-tagged C220 (G-C220) and C220 mutations for nuclear exporting signal sequence (NES) (G-C220mNES). The left and middle fluorescence images were acquired from the same field of view using either high or low gain of the detector. Bar = 5 µm. (B) Dots and bar graphs indicate relative fluorescence intensity in the nucleoplasm to that in the cytoplasm (N/C ratio). (C) Dots and bar graphs indicate the population of the cells containing cytoplasmic condensates (mean ± 95% CI; n = 3). ND: Not determined; *p*-values above the lines: one-way ANOVA with Tukey's test.

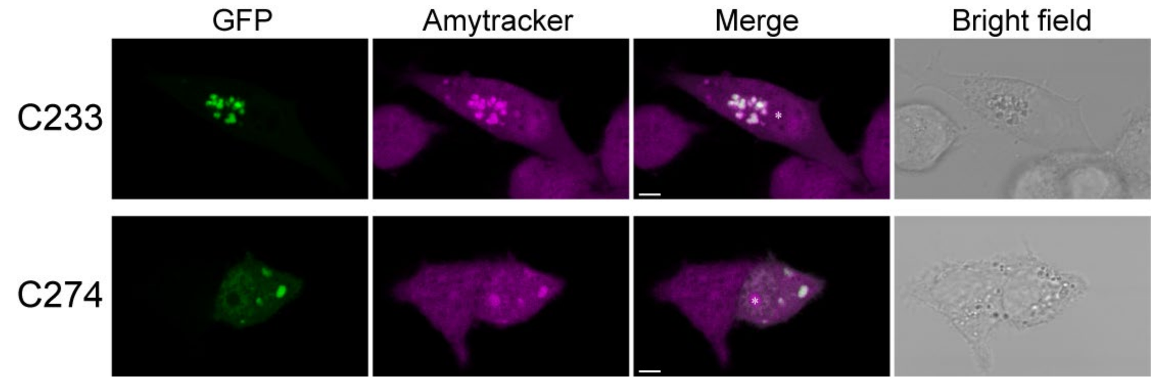


**Supplemental Figure 2: Amytracker staining of the C233 and C274 condensates in Neuro2a cells**
Confocal images of GFP-tagged C233 and C274-expressing Neuro2a cells stained with a fluorescent tracer for amyloid, Amytracker. The asterisk in the images represents the nucleolus. Bars = 5 μm.

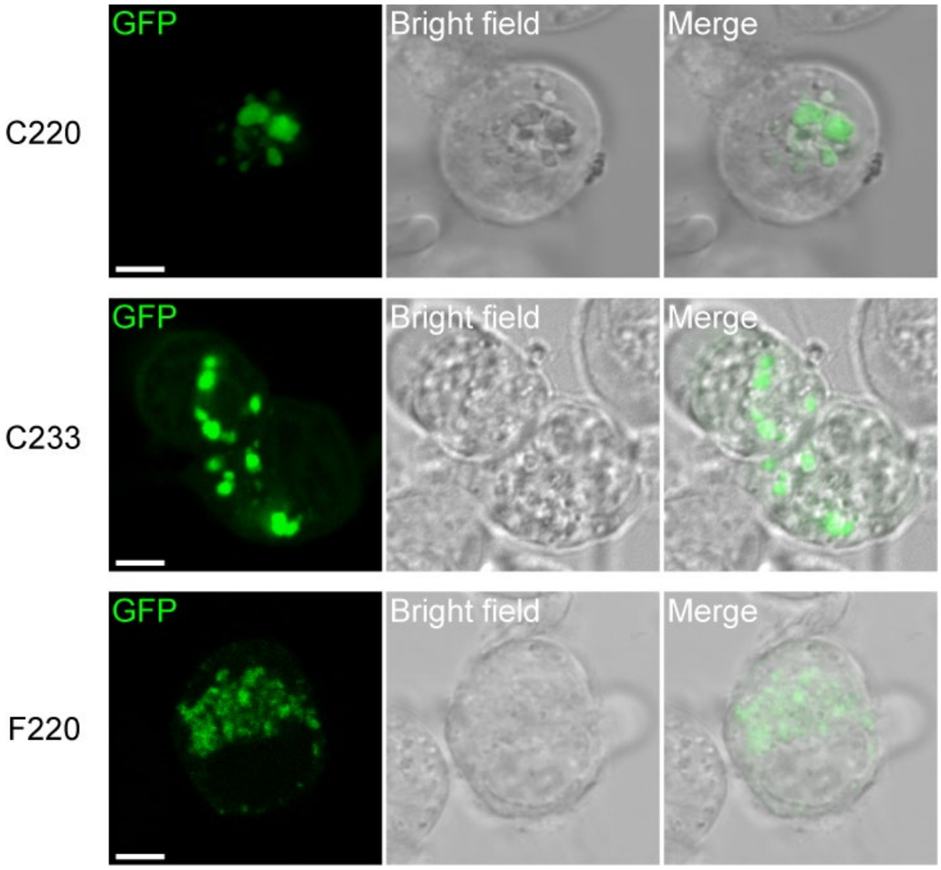


**Supplemental Figure 3: Confocal fluorescent images of Neuro2a cells expressing C220, C233, and F220 tagged with GFP after the 1,6-Hexanediol treatment.**

The confocal fluorescence images of cells containing cytoplasmic condensates were treated with 1,6-hexanediol were represented. Bars = 5 μm.

A

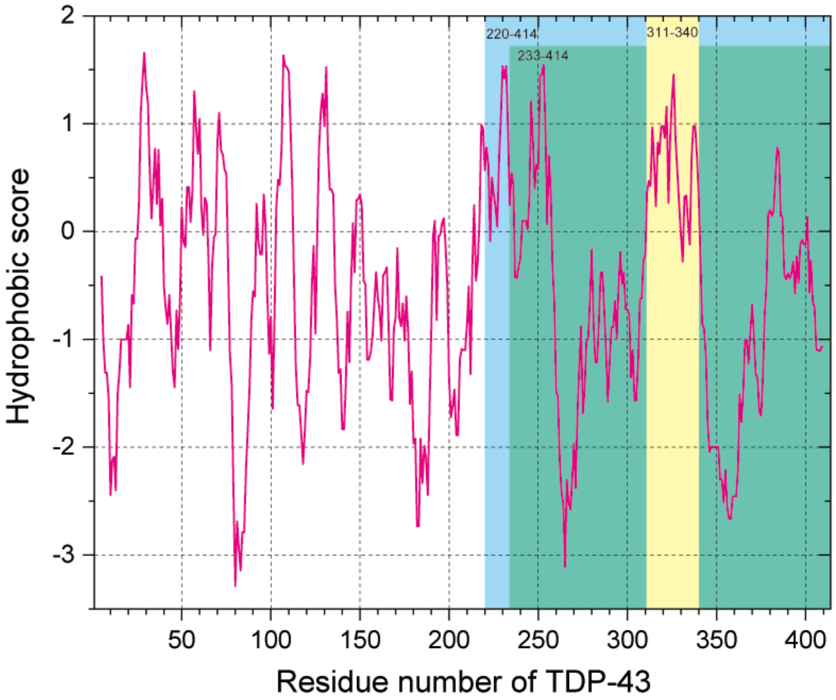


B

| Amino acid | Score |
|---|---|
| Ile | 4.5 |
| Val | 4.2 |
| Leu | 3.8 |
| Phe | 2.8 |
| Cys | 2.5 |
| Met | 1.9 |
| Ala | 1.8 |
| Gly | -0.4 |
| Thr | -0.7 |
| Ser | -0.8 |
| Trp | -0.9 |
| Tyr | -1.3 |
| Pro | -1.6 |
| His | -3.2 |
| Asn | -3.5 |
| Asp | -3.5 |
| Gln | -3.5 |
| Glu | -3.5 |
| Lys | -3.9 |
| Arg | -4.5 |

**Supplemental Figure 4: Hydropathy plot of human TDP-43 amino acid sequence**

(A) The hydropathy plot was calculated using the Expasy Web server [Kyte J. & Doolittle R.F., ***J. Mol. Biol.***, 157, 105-132 (1982)]. The CTF regions of C220 and C233 represented a cyan and green area, respectively. A highly hydrophobic region in GRR was shown in a yellow area. (B) The hydrophobicity scores of each amino acid are represented (Red to blue: high to low hydrophobicity).

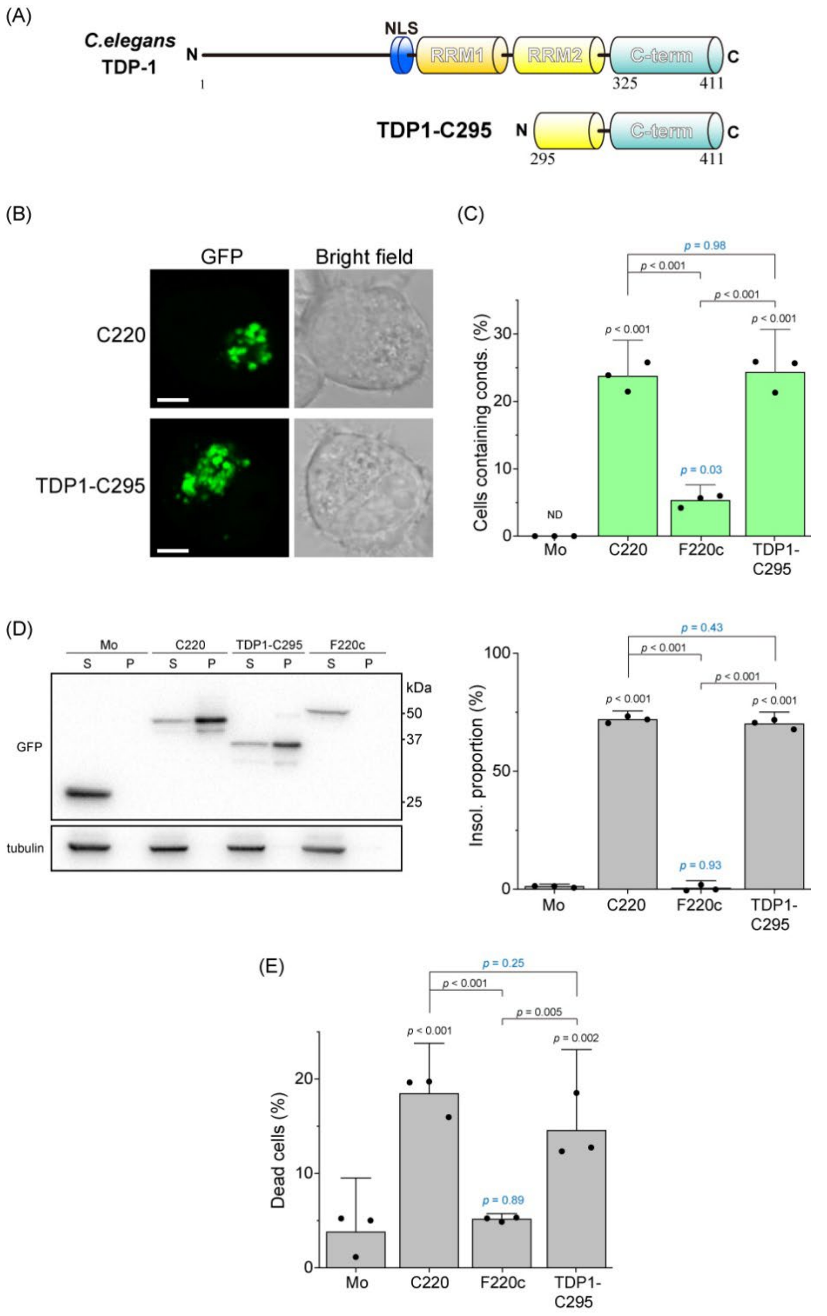


**Supplemental Figure 5 Formation of the cytoplasmic condensates and toxicity of *Caenorhabditis elegans* TDP-1 carboxy terminal fragment in Neuro2a cells**

(A) Primary structures and abbreviation of *Caenorhabditis elegans* TDP-1 and its carboxy-terminal fragment (TDP1-C295: 295–411 amino acids) that corresponds to human C220/TDP25. The RRM and C-term denote the RNA/DNA-recognition motif and the C-terminal region, respectively. (B) Confocal fluorescence and bright field microscopic images of Neuro2a cells expressing GFP-tagged C220 and TDP1-C295. Bars = 5 μm. (C) The population of Neuro2a cells containing cytoplasmic condensates of GFP-tagged C220, tRRM2 (F220c), TDP1-C295, and GFP monomers (Mo). Bars: mean + 95% CI; Dots: raw values (the number of trials (n) = 3; the number of counted cells per a trial (N) > 200). (D) Western blot stained using an anti-GFP and anti-α-tubulin antibody of cell lysates: 1% SDS-soluble

(S) and insoluble (P) (*left*). The quantification of the abundance of TDP-43 CTFs, TDP1-C295, and GFP monomers (Mo) in the insoluble fraction. The abundance shows the normalized band intensity in the P fraction to total (S+P) fraction (n = 3) (bottom). (E) Population of dead cells expressing TDP-43 CTFs and TDP1-C295 tagged with GFP and GFP monomers (Mo) using a propidium iodide exclusion test. Bars: mean and 95% CI (n = 3; N > 200). (C, D, and E) *p*-values above the bars and lines: one-way ANOVA with Tukey's test compared to GFP monomers as control and comparison between lines, respectively. ND: not determined.

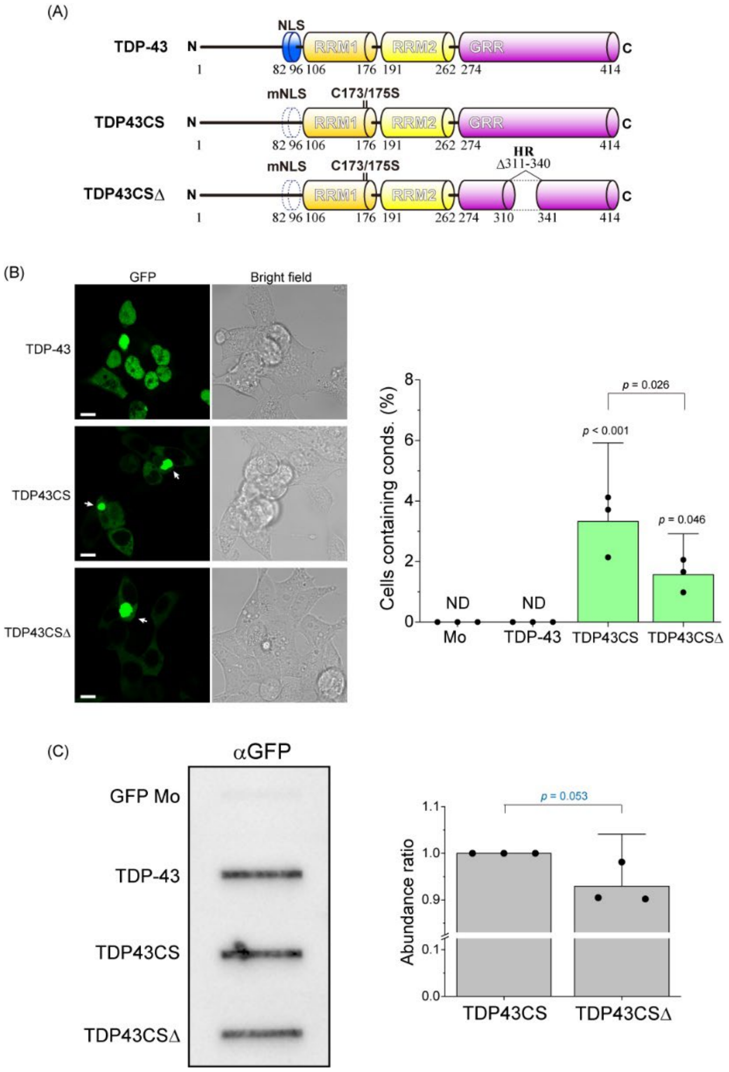


**Supplemental Figure 6 Formation of the cytoplasmic condensates of full-length TDP-43 in Neuro2a cells**

(A) Primary structures and abbreviation of human TDP-43 and its mutants. The RRM and GRR denote the RNA/DNA-recognition motif and the glycine-rich region, respectively. TDP43CS carries mutations in nuclear localization signal sequence (mNLS) and two substitutions of 173$^{th}$ and 175$^{th}$ cysteine to serine residues. TDP43CSΔ lacks hydrophobic region (HR) in GRR. All proteins are tagged with monomeric GFP at the C-terminus. (B) Confocal fluorescence and bright field microscopic images of Neuro2a cells expressing GFP-tagged TDP-43, TDP43CS, and TDP43CSΔ. Bars = 10 μm. (C) The population of Neuro2a cells containing cytoplasmic condensates of GFP-tagged TDP-43, TDP43CS, TDP43CSΔ, and GFP monomers (Mo). Bars: mean + 95% CI; Dots: raw values (the number of trials (n) = 3; the number of counted cells per a trial (N) > 200). (D) Filter retardation assay followed by staining using an anti-GFP antibody of cell lysates (*left*). The quantification of the abundance ratio of TDP43CS and TDP43CSΔ retarded on the membrane (n = 3) (*right*). *p*-values above the bars and lines: one-way ANOVA with Tukey's test compared to GFP monomers as control and comparison between lines, respectively. ND: not determined.

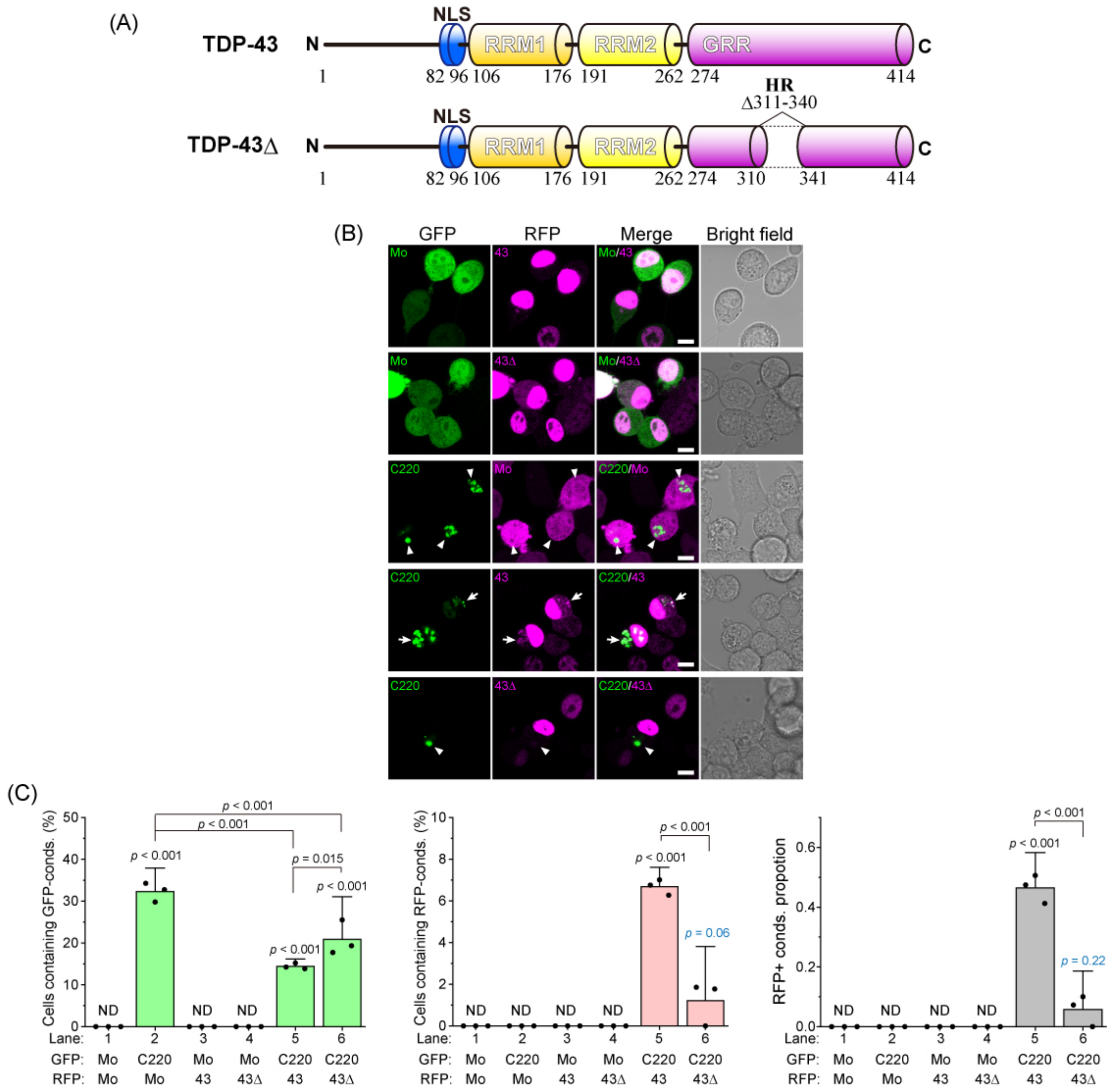


**Supplemental Figure 7 The role of hydrophobic region in TDP-43 glycine-rich region during co-condensation between C220 and full-length TDP-43.**

(A) Domain structure and abbreviation of human full-length TDP-43 and that lacking 311-340 region (TDP43Δ). HR and GRR denote a hydrophobic region and a glycine-rich region, respectively. (B) Confocal fluorescence and bright field microscopic images of Neuro2a cells expressing GFP-tagged C220 and RFP-tagged TDP-43 (43) or TDP43Δ (43Δ). Mo indicates GFP or RFP monomers. White arrowheads represent the position of green fluorescence-positive but red fluorescence-negative condensates in the cytoplasm. White arrows represent both green and red fluorescence-positive condensates in the cytoplasm. Bars = 5 μm. (C) *left*: The population of Neuro2a cells that contain GFP-positive condensates (conds) in the cytoplasm. *Middle*: The population of Neuro2a cells containing RFP-positive condensates in the cytoplasm. *Right*: The ratio of RFP-positive condensates to GFP-positive ones. ND: Not determined; N/A: Not applicable; *p*-values above the lines: one-way ANOVA with Tukey's test (the number of trials (n) = 3; the number of counted cells per trial (N) > 300).

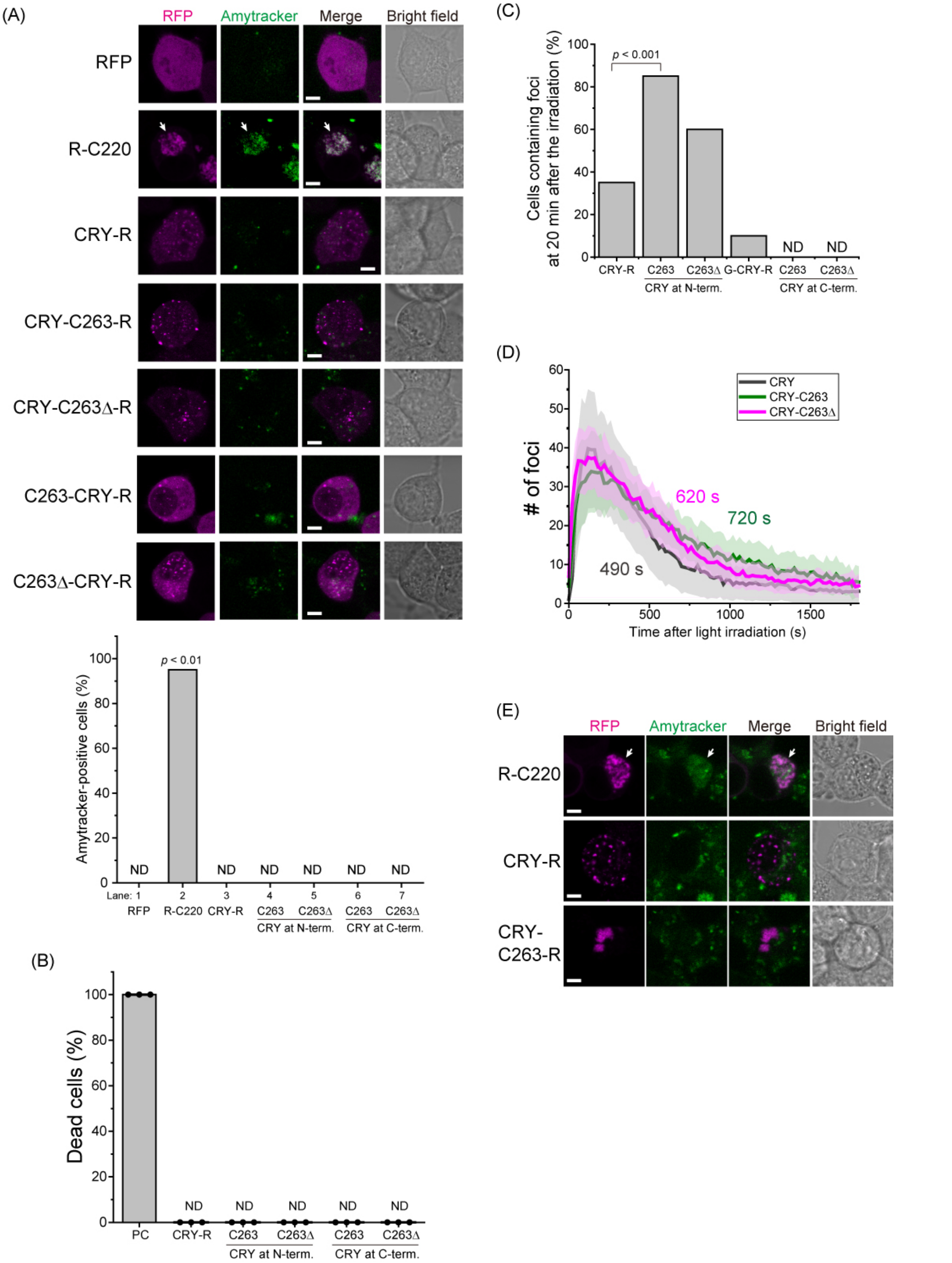


**Supplemental Figure 8: Fate and consequence of light-induced condensates of TDP-43 carboxy-terminal fragments.**

(A) *Top*: Fluorescence images of Amytracker 520-stained Neuro2a cells expressing TDP-43 CTFs: C220, C220Δ, C263, and C263Δ tagged with a cryptochrome-derived oligomerization inducer, CRY2olig (CRY), RFP monomers (RFP), and RFP-tagged C220 at 30 min after the 488 nm-light irradiation. The order of the hyphens before and after indicates the N/C-terminal side of the CRY tag. White arrowheads represent Amytracker-positive condensates. Bars = 5 μm. *Bottom*: Population of cells that form foci after the blue-light irradiation obtained from all measured cells. ND: Not determined; *p*-values above the bars: hypothesis test for the difference in the population proportions compared to RFP monomers. (B) Population of dead cells using a DRAQ7 dye exclusion test. PC denotes ethanol-treated cells as a positive control. Bars: mean and 95% CI (the number of trials (n) = 3; the number of counted cells per trial (N) > 300). (C) Population of cells harboring foci at 20 min after the light irradiation. *p*-values were obtained by hypothesis test for the difference in the population proportions compared to CRY-R. (D) Time-course of the number of light-induced foci of CRY, CRY-C263, and CRY-C263Δ in a cell after turning off the blue light (dark gray, green, and magenta, respectively). The thick solid line represents the mean, and the shaded area in a lighter color indicates ± 95% CI (the number of cells = 20). Inset numbers indicate mean half-decreasing times after their maximum peak. (E) Amytracker-stained Neuro2a cells harboring TDP25, CRY, and CRY-C263 foci that remained after 8 hours of blue light irradiation. White arrowheads indicate Amytracker-positive foci. Bars = 5 μm.

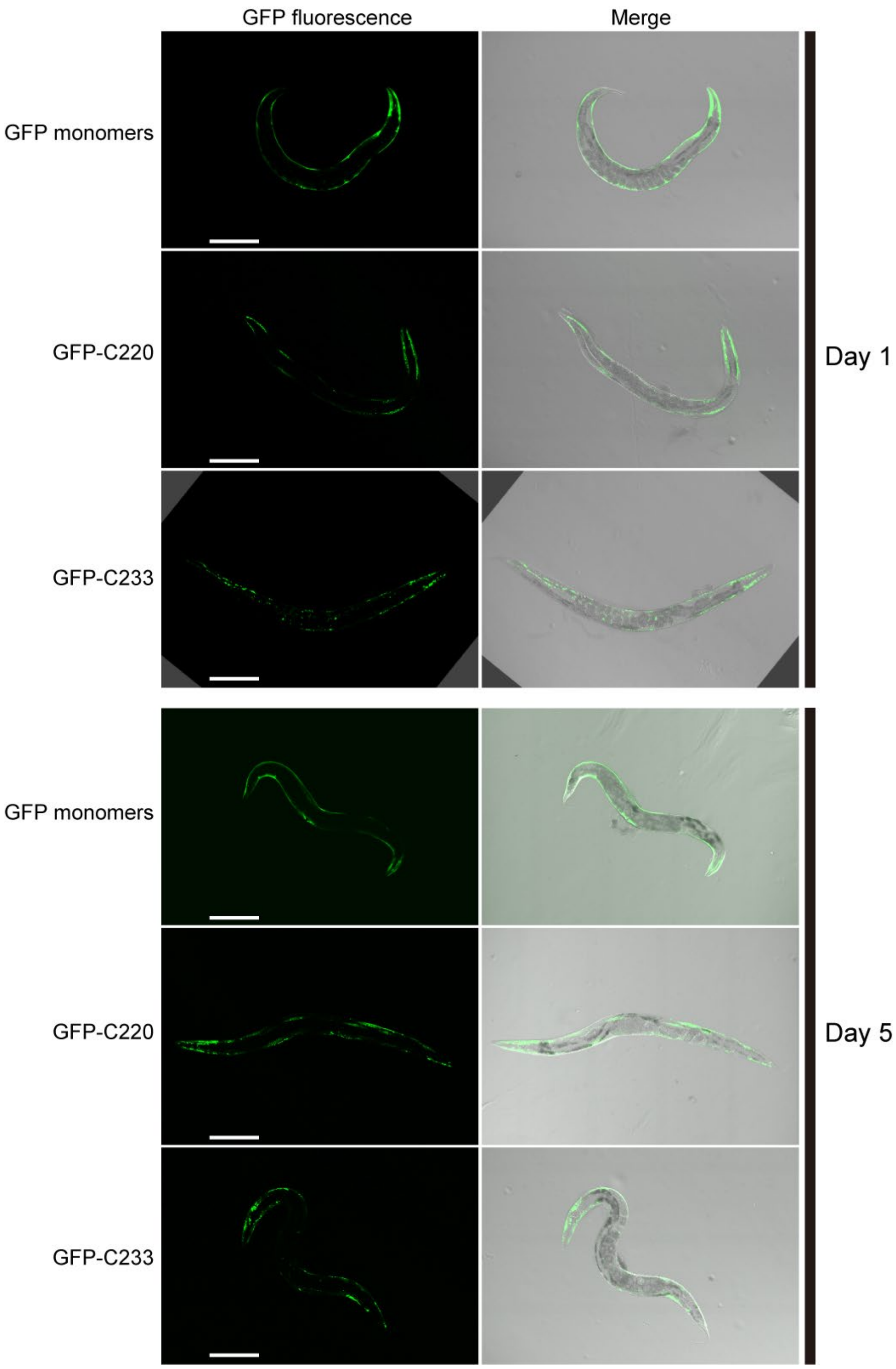


**Supplemental Figure 9: Confocal fluorescence images of nematodes expressing GFP, GFP-C220, and GFP-C233.**

Total view of the nematodes expressing GFP-C220, GFP-C233, and GFP monomers on 4 or 8 days after synchronization by bleaching (Day 1 and 5 of adulthood, respectively) shown in Figure 8A. Bars = 0.2 mm.

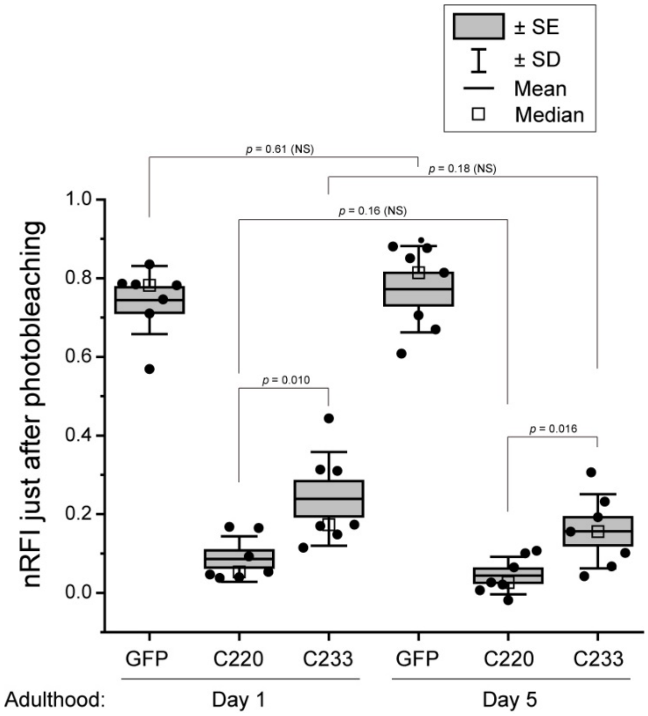


**Supplemental Figure 10: Comparison of normalized relative fluorescence intensities (nRFI) just after photobleaching of C220 and C233 tagged with GFP in *C. elegans*.**

Box and bars with dots plot of nRFI of GFP monomers, GFP-C220, and GFP-C233 just after photobleaching (t = 0 s in Figure 9A). *p*-values above the lines: one-way ANOVA with Tukey's test (n =7); NS indicates $p > 0.05$.

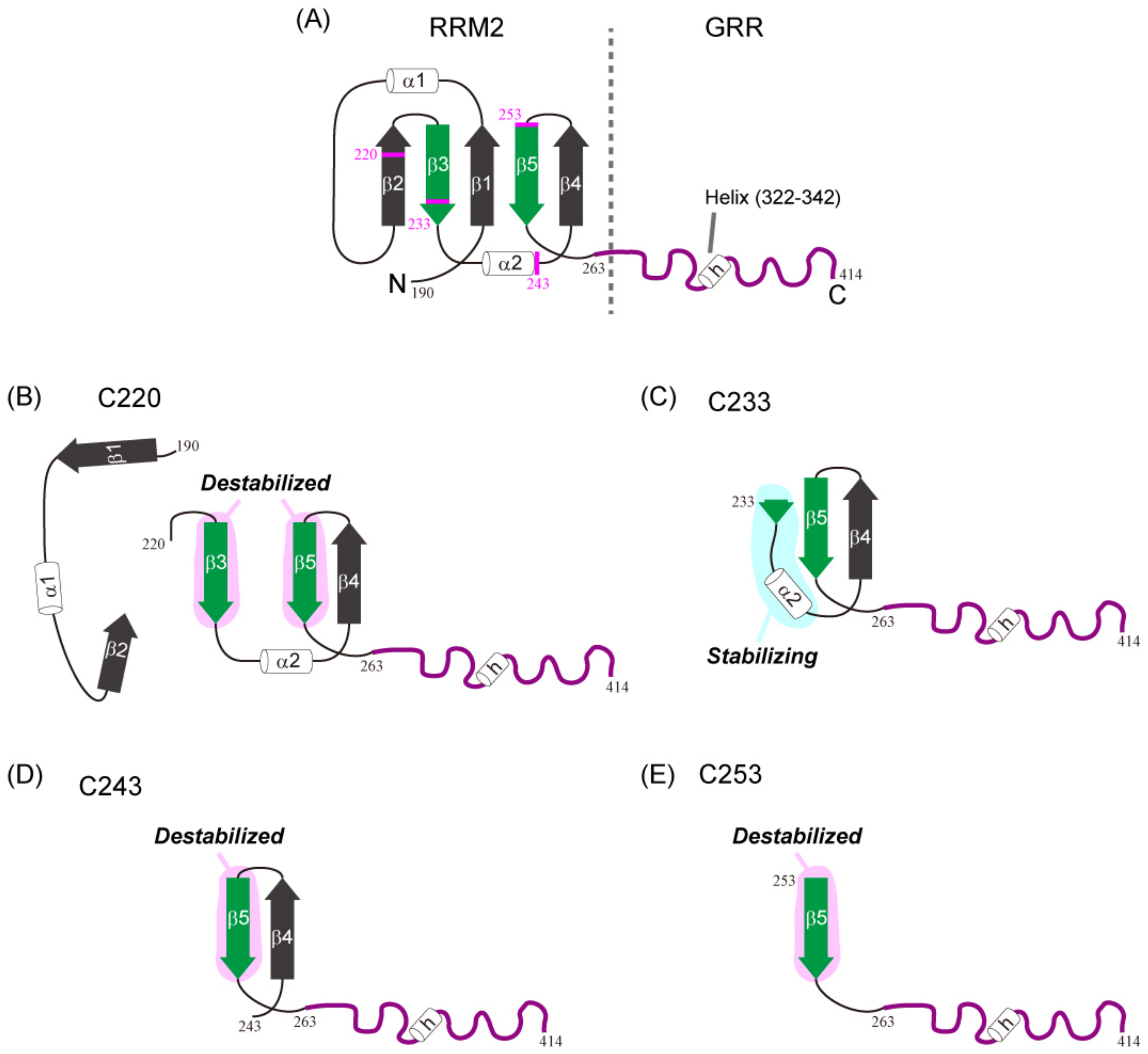


**Supplemental Figure 11: Illustration of the secondary structure of intact RRM2 and GRR of TDP-43 and the cleavage positions of C220, C233, C243, and C253**

(A–E) α1 and α2 indicate two α-helixes and β1–5 indicate five β-strands in RRM2 of TDP-43 observed in PDB#3D2W. The h means α-helical structure in GRR of TDP-43 observed in PDB#2N3X [1]. Structure of intact RRM2 and GRR (A), C220/TDP25 and the cleaved fragment of RRM2 (B), C233 (C), C243 (D), and C253 (E).

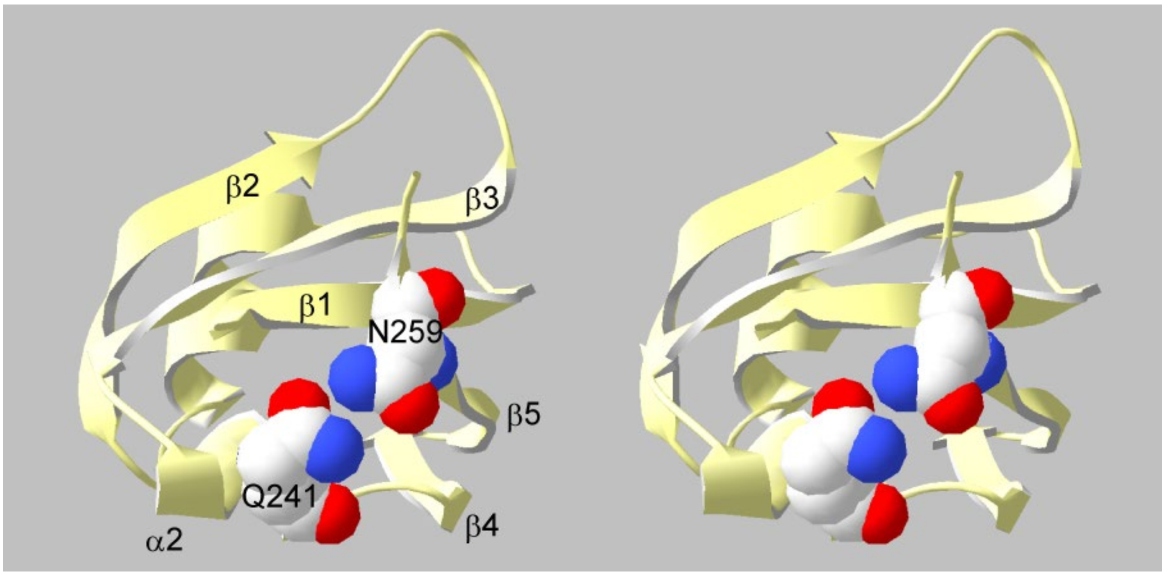


**Supplemental Figure 12: Stereoview of the electrostatic interaction of the amino acid side chain (Q241 and N259) in RRM2**

α1 and α2 indicate two α-helixes and β1–5 indicate five β-strands in RRM2 of TDP-43 observed in PDB#3D2W. The balls indicate the atoms of the side chain of Q241 and N259 (red and blue balls represent atoms having positive and negative charges, respectively).
